# A Variational Inference Approach to Single-Cell Gene Regulatory Network Inference using Probabilistic Matrix Factorization

**DOI:** 10.1101/2022.09.09.507305

**Authors:** Omar Mahmood, Claudia Skok Gibbs, Richard Bonneau, Kyunghyun Cho

## Abstract

Inferring gene regulatory networks (GRNs) from single cell data is challenging due to heuristic limitations and a lack of uncertainty estimates in existing methods. To address this, we present Probabilistic Matrix Factorization for Gene Regulatory Network Inference (PMF-GRN). Using single cell expression data, PMF-GRN infers latent factors capturing transcription factor activity and regulatory relationships, incorporating experimental evidence via prior distributions. By utilizing variational inference, we facilitate hyperparameter search for principled model selection and direct comparison to other generative models. We extensively test and benchmark our method using single cell datasets from *Saccharomyces cerevisiae*, human Peripheral Blood Mononuclear Cells (PBMCs), and BEELINE synthetic data. We discover that PMF-GRN infers GRNs more accurately than current state-of-the-art single-cell GRN inference methods, offering well-calibrated uncertainty estimates for additional interpretability.

## Background

An essential problem in systems biology is to extract information from genome wide 5 sequencing data to unravel the mechanisms controlling cellular processes within 6 heterogeneous populations [1]. Gene regulatory networks (GRNs) that annotate 7 regulatory relationships between transcription factors (TFs) and their target genes [2] have proven to be useful models for stratifying functional differences between cells 9 [3, 4, 5, 6] that can arise during normal development [7], responses to environmental 10 signals [8] and dysregulation in the context of disease [9, 10, 11].

GRNs cannot be directly measured with current sequencing technology. Instead, methods must be developed to piece together snapshots of transcriptional processes 13 in order to reconstruct a cell’s regulatory landscape [12]. Initial approaches to GRN 14 inference relied on Microarray technology [13, 14, 15], a hybridization-based method 15 to measure the expression of thousands of genes simultaneously [16]. This technology 16 was biased as it was limited to only those genes that were annotated at the time, 17 which in turn presented challenges for inferring the complete regulatory landscape [1]. 18 Subsequently, the high-throughput sequencing method RNA-seq provided a genome 19 wide readout of transcriptional output, allowing for the detection of novel transcripts [17] and thus improving GRN inference potential. More recently, single-cell RNA-seq technology has enabled the characterization of gene expression profiles within heterogeneous populations [18], vastly increasing the potential for GRN inference algorithms [19, 20]. In contrast to bulk RNA experiments (Microarray and RNA-seq) that average measurements of gene expression across heterogenous cell populations, GRNs inferred from single-cell data have the advantage of unmasking biological signal in individual cells [21].

Several matrix factorization approaches have been proposed to overcome the limitations of reconstructing GRNs from Microarray data [22]. These include use of statistical techniques such as Singular Value Decomposition and Principal Component Analysis [23], Bayesian Decomposition [24], and Non-negative Matrix Factorization [25, 26, 27]. More recently, matrix factorization approaches have been applied to integrative analysis of DNA methylation and miRNA expression data [28], as well as single-cell RNA-seq and single-cell ATAC-seq data [29]. However, to the best of our knowledge, these matrix factorization approaches have not yet been used to infer GRNs from single-cell gene expression data. Meanwhile, several regression-based methods have been proposed to learn GRNs from single-cell RNA-seq and single-cell ATAC-seq to capture regulatory relationships at single-cell resolution [30]. So far, these integrative approaches to GRN inference have been successfully implemented using regularized regression [31], self-organizing maps [32], tree-based regression [33], and Bayesian Ridge regression [34].

Although regression-based methods for inferring GRNs from single-cell data are available, they still suffer from significant limitations [35]. Firstly, these methods are designed for specific input datasets, such as bulk or single-cell RNA-seq, causing issues when new data becomes available or new assumptions are required in the model. This can result in inaccurate predictions if the new data or assumptions are not well integrated into the existing model, leading to the need for a complete re-design of the algorithm, which can be costly and time-consuming. Additionally, these methods typically focus on inferring a single GRN that explains the available data, without performing hyperparameter search to determine the optimal model. This can lead to heuristic model selection, with no justification for the approach taken or evidence that the best possible model has been selected. Conversely, hyperparameter search ensures the accuracy of the GRN inference algorithm by finding the optimal model that fits the data well while avoiding overfitting. Regression-based GRN inference algorithms that do not perform hyperparameter search may miss important data features or overemphasize irrelevant ones, leading to inaccurate or incomplete models. Moreover, these methods do not provide an indication of their uncertainty about the predictions that they make. Finally, several regression-based GRN inference algorithms struggle to scale optimally to the size of typical single-cell datasets, limiting inference to small subsets of data or requiring enormous amounts of computational time.”

In this study, we introduce PMF-GRN, a novel approach that uses probabilistic matrix factorization [36] to infer gene regulatory networks from single-cell gene expression and chromatin accessibility information. This approach extends previous methods that applied matrix factorization for GRN inference with Microarray data, to address the current limitations in regression-based single-cell GRN inference. We implement our approach in a probabilistic setting with variational inference, which provides a flexible framework to incorporate new assumptions or biological data as required, without changing the way the GRN is inferred. We also use a principled hyperparameter selection process, which optimizes the parameters of our probabilistic model for automatic model selection. In this way, we replace heuristic model selection by comparing a variety of generative models and hyperparameter configurations before selecting the optimal parameters with which to infer a final GRN. Our probabilistic approach provides uncertainty estimates for each predicted regulatory interaction, serving as a proxy for the model confidence in each predicted interaction. Uncertainty estimates can be useful in the situation where there are limited validated interactions or a gold standard is incomplete. By using stochastic gradient descent (SGD), we perform GRN inference on a GPU, allowing us to easily scale to a large number of observations in a typical single-cell gene expression dataset. Unlike many existing methods, PMF-GRN is not limited by pre-defined organism restrictions, making it widely applicable for GRN inference.

To demonstrate the novelty and advantages of PMF-GRN, we apply our method to datasets from *Sacchromyces cerevisiae*, human Peripheral Blood Mononuclear Cells (PBMCs) and BEELINE. In our first experiment, we apply our method to two single cell gene expression datasets for the model organism *S. cerevisiae*. We evaluate our model’s performance in a normal inference setting, as well as with cross-validation and noisy data. To assess the accuracy of predicted regulatory interactions, we evaluate all regulatory predictions using Area Under the Precision Recall Curve (AUPRC) against database derived gold standards. Our findings show that the uncertainty estimates are well-calibrated for inferred TF-target gene interactions, as the accuracy of predictions increases when the associated uncertainty decreases. Here, in comparison to three state-of-the-art regression-based methods for inferring single cell GRNs, namely the Inferelator [31], Scenic [33], and Cell Oracle [34], our method demonstrates an overall improved performance in recovering the true underlying GRN. Additionally, we apply our method to a PBMC dataset and explore the inferred TFA profiles in the context of annotated cell types and specific immune TFs. We investigate regulatory edges in our inferred GRN and find compelling support for our predictions. Lastly, we benchmark our method using six synthetic datasets generated from BEELINE [37] and demonstrate consistent outperformance of PMF-GRN compared to the baseline.

## Results

### The PMF-GRN Model

The goal of our probabilistic matrix factorization approach is to decompose observed gene expression into latent factors, representing TF activity (TFA) and regulatory interactions between TFs and their target genes. These latent factors, which represent the underlying GRN, cannot be measured experimentally, unlike gene expression. We model an observed gene expression matrix *W* ∈ ℝ^N ×M^ using a TFA matrix 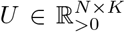, a TF-target gene interaction matrix *V* ∈ ℝ^M ×K^, observation noise σobs ∈ (0, ∞) and sequencing depth d ∈ (0, 1)N, where *N* is the number of cells, *M* is the number of genes and *K* is the number of TFs. We rewrite *V* as the product of a matrix *A* ∈ (0, 1)^M×K^, representing the degree of existence of an interaction, and a matrix *B* ∈ ℝ^M ×K^ representing the interaction strength and its direction:

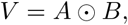

where ⊙ denotes element-wise multiplication. An overview of the graphical model is shown in Figure 1A.

**Figure 1:**
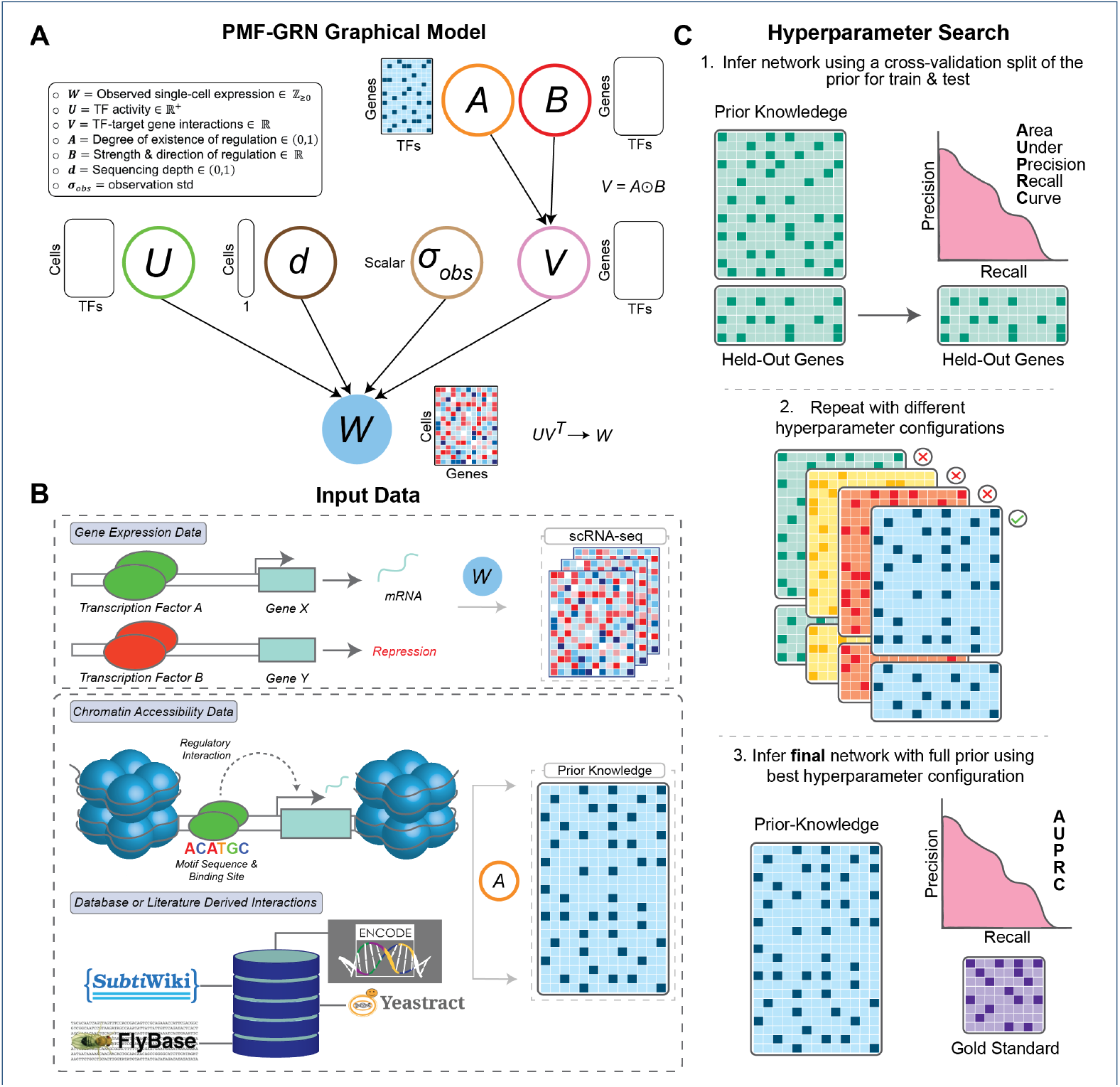
(**A**) PMF-GRN graphical model overview. Input single-cell gene expression *W* is decomposed into several latent factors. Information obtained from chromatin accessibility data or genomics databases is incorporated into the prior distribution for *A*. (**B**) Input experimental data for PMF-GRN includes single-cell RNA-seq gene expression data. Prior-known TF-target gene interactions can be obtained using chromatin accessibility in parallel with known TF motifs, or through databases or literature derived interactions.(**C**) Hyperparameter selection process is performed for optimal model selection. The provided prior-known network is split into a train and validation dataset. 80% of the prior-known information is used to infer a GRN, while the remaining 20% is used for validation by computing AUPRC. This process is repeated multiple times, using different hyperparameter configurations in order to determine the optimal hyperparameters for the GRN inference task at hand. Finally, using the optimal hyperparameters, a final network is inferred using the full prior and evaluated using an independent gold standard.

These latent variables are mutually independent a priori, i.e., *p*(*U, A, B, σ*_*obs*_, *d*) *= p*(*U* )*p*(*A*)*p*(*B*)*p*(*σ*_*−−*_The observations W result from a matrix product *UV* ^⊤^. We assume noisy observations by defining a distribution over the observations with the level of noise σobs, i.e., *p*(*W* |*U, V* = *A* ⊙ *B, σ*_*obs*_, *d*).

For the matrix *A*, prior hyperparameters represent an initial guess of the interaction between each TF and target gene which need to be provided by a user. These can be derived from genomic databases or obtained by analyzing other data types, such as the measurement of chromosomal accessibility, TF motif databases, and direct measurement of TF-binding along the chromosome, as shown in Figure 1B (see Methods section for details).

Given this generative model, we perform posterior inference over all the unobserved latent variables; *U, A, B, d* and *σ*_*obs*_, and use the posterior over A to investigate TFtarget gene interactions. Exact posterior inference with an arbitrary choice of prior and observation probability distributions is, however, intractable. We address this issue by using variational inference [38, 39], where we approximate the true posterior distributions with tractable, approximate (variational) posterior distributions.

We minimize the KL-divergence *D*_*KL*_(*q*∥*p*) between the two distributions with respect to the parameters of the variational distribution *q*, where *p* is the true posterior distribution. This allows us to find an approximate posterior distribution *q* that closely resembles *p*. This is equivalent to maximizing the evidence lower bound (ELBO) i.e. a lower bound to the marginal log likelihood of the observations W :

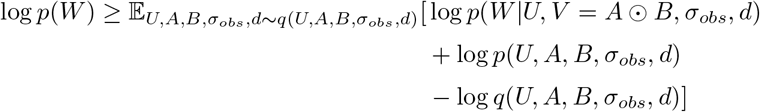

The mean and variance of the approximate posterior over each entry of A, obtained from maximizing the ELBO, are then used as the degree of existence of an interaction between a TF and a target gene and its uncertainty, respectively.

It is important to note that matrix factorization based GRN inference is only identifiable up to a latent factor (column) permutation. In the absence of prior information, the probability that the user assigns TF names to the columns of *U* and *V* in the same order that the inference algorithm implicitly assigns TFs to these columns is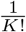, is essentially 0 for any reasonable value of *K*. Incorporating prior-knowledge of TF-target gene interactions into the prior distribution over *A* is therefore essential in order to provide the inference algorithm with the information of which column corresponds to which TF.

With this identifiability issue in mind, we design an inference procedure that can be used on any prior-knowledge edge matrices, described in Figure 1C. The first step is to randomly hold out prior information for some percentage of the genes in *p(A)* (we choose 20%) by leaving the rows corresponding to these genes in A but setting the prior logistic normal means for all entries in these rows to be the same low number.

The second step is to carry out a hyperparameter search using this modified prior-knowledge matrix. The early stopping and model selection criteria are both the ‘validation’ AUPRC of the posterior point estimates of A, corresponding to the held out genes, against the entries for these genes in the full prior hyperparameter matrix. This step is motivated by the idea that inference using the selected hyperparameter configuration should yield a GRN whose columns correspond to the TF names that the user has assigned to these columns.

The third step is to choose the hyperparameter configuration corresponding to the highest validation AUPRC and perform inference using this configuration with the full prior. An importance weighted estimate of the marginal log likelihood is used as the early stopping criterion for this step. The resulting approximate posterior provides the final posterior estimate of *A*.

### Advantages of PMF-GRN

Existing methods almost always couple the description of the data generating process with the inference procedure used to obtain the final estimated GRN [31, 34, 33]. Designing a new model thus requires designing a new inference procedure specifically for that model, which makes it difficult to compare results across different models due to the discrepancies in their associated inference algorithms. Furthermore, this *ad hoc* nature of model building and inference algorithm design often leads to the lack of a coherent objective function that can be used for proper hyperparameter search as well as model selection and comparison, as evident in [31]. Heuristic model selection in available GRN inference methods presents the challenge of determining and selecting the optimal model in a given setting.

The proposed PMF-GRN framework decouples the generative model from the inference procedure. Instead of requiring a new inference procedure for each generative model, it enables a single inference procedure through (stochastic) gradient descent with the ELBO objective function, across a diverse set of generative models. Inference can easily be performed in the same way for each model. Through this framework, it is possible to define the prior and likelihood distributions as desired with the following mild restrictions: we must be able to evaluate the joint distribution of the observations and the latent variables, the variational distribution and the gradient of the log of the variational distribution.

The use of stochastic gradient descent in variational inference comes with a significant computational advantage. As each step of inference can be done with a small subset of observations, we can run GRN inference on a very large dataset without any constraint on the number of observations. This procedure is further sped up by using modern hardware, such as GPUs.

Under this probabilistic framework, we carry out model selection, such as choosing distributions and their corresponding hyperparameters, in a principled and unified way. Hyperparameters can be tuned with regard to a predefined objective, such as the marginal likelihood of the data or the posterior predictive probability of held out parts of the observations. We can further compare and choose the best generative model using the same procedure.

This framework allows us to encode any prior knowledge via the prior distributions of latent variables. For instance, we incorporate prior knowledge about TF-gene interactions as hyperparameters that govern the prior distribution over the matrix *A*. If prior knowledge about TFA is available, this can be similarly incorporated into the model via the hyperparameters of the prior distribution over *U* .

Because our approach is probabilistic by construction, inference also estimates uncertainty without any separate external mechanism. These uncertainty estimates can be used to assess the reliability of the predictions, i.e., more trust can be placed in interactions that are associated with less uncertainty. We verify this correlation between the degree of uncertainty and the accuracy of interactions in the experiments. Overall, the proposed approach of probabilistic matrix factorization for GRN inference is scalable, generalizable and aware of uncertainty, which makes its use much more advantageous compared to most existing methods.

### PMF-GRN Recovers True Interactions in Simple Eukaryotes

To evaluate PMF-GRN’s ability to infer informative and robust GRNs, we leverage two single-cell RNA-seq datasets from the model organism *Saccharomyces cerevisiae* [8, 40]. This eukaryote, being relatively simple and extensively studied, provides a reliable gold standard [41] for assessing the performance of different GRN inference methods. We conduct three experiments to compare the performance of three stateof-the-art GRN inference methods, the Inferelator (AMuSR, BBSR, and StARS) [31], SCENIC [33], and CellOracle [34]. Throughout these experiments, each method is provided with the exact same single-cell RNA-seq datasets (GSE125162 [8]: N cells = 38, 225, GSE144820 [40]: N cells = 6, 118, combined: N cells = 44, 343 by M genes = 6, 763), prior-knowledge (M genes = 6, 885 by K TFs = 220) and gold standard (M genes = 993 by K TFs = 98).

In the first experiment, we infer GRNs for each of the two yeast datasets and average the posterior means of *A* to simulate a “multi-task” GRN inference approach. Using AUPRC, we demonstrate that PMF-GRN outperforms AMuSR, StARS, and SCENIC, while performing competitively with BBSR and CellOracle (Figure 2A). We next combine the two expression datasets into one observation to test whether each method can discern the overall GRN accurately when data is not cleanly organized into tasks. This experiment reveals a substantial performance decrease for BBSR, indicating its dependence on organized gene expression tasks. This finding suggests potential challenges for BBSR in more complex organisms with less well-defined cell types or conditions. For benchmarking purposes we provide two negative controls for each method, a GRN inferred without prior information (No Prior), and a GRN inferred using shuffled prior information (Shuffled Prior). For all methods, these negative controls achieve an expected low AUPRC. It is essential to note that for CellOracle, an experiment with no prior information could not be performed. This is due to the fact that by design, CellOracle cannot learn regulatory edges that are not included in the prior information.

**Figure 2:**
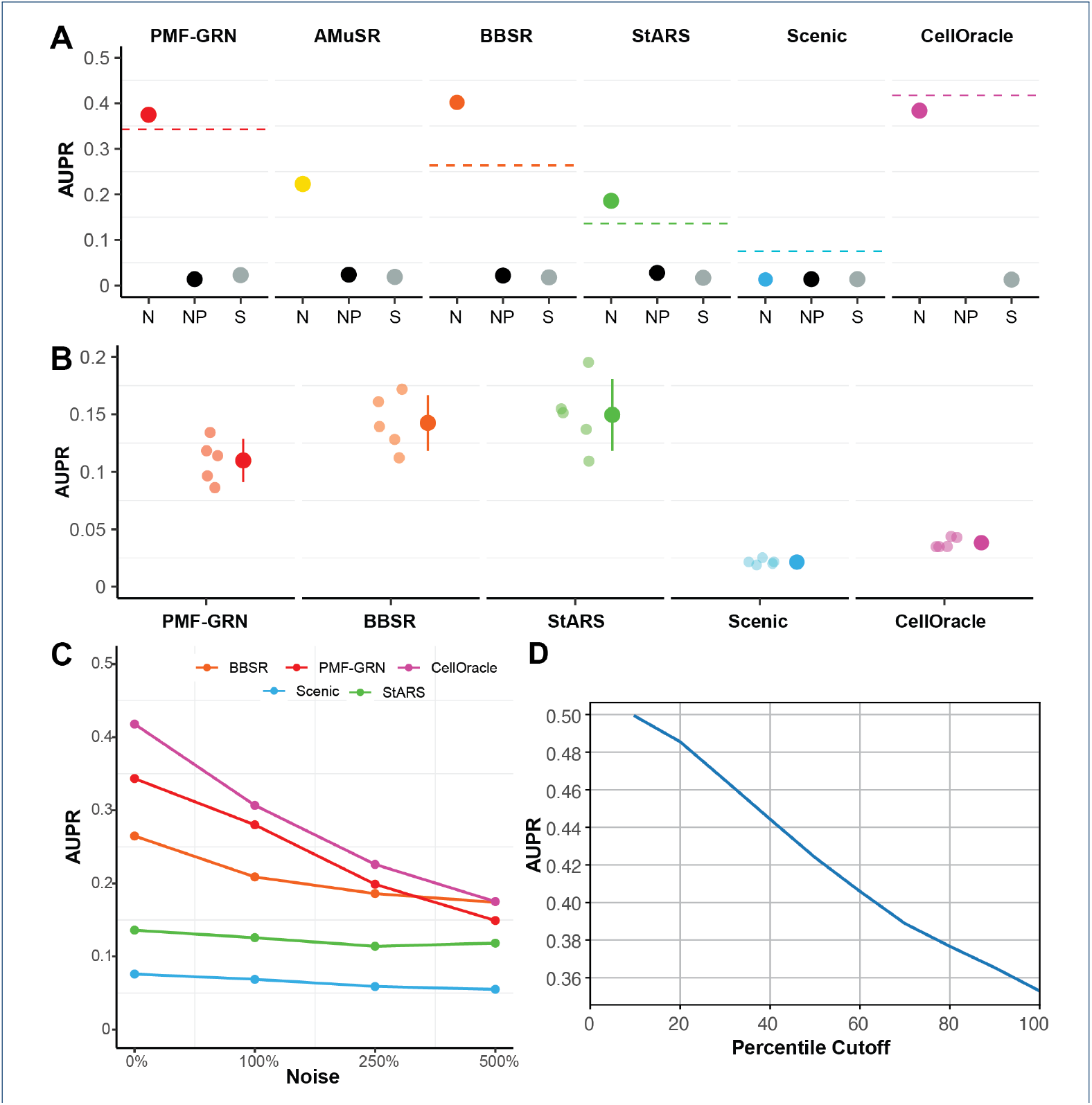
GRN inference in *S. cerevisiae*. (**A**) Consensus Network AUPR with a normal prior-knowledge matrix (N): PMF-GRN (red) performance compared to Inferelator algorithms (AMuSR in yellow, BBSR in orange, StARS in green), SCENIC (blue), and CellOracle (purple). Dashed line represents the baseline if expression data is combined. Negative controls: no prior information (NP - black) and shuffled prior information (S - gray). (**B**) 5-Fold Cross-Validation Baseline: Each dot with low opacity represents one of the five experiments. Colored dots and lines depict the mean AUPR *±* standard deviation for each GRN inference method. (**C**) GRNs inferred with increasing amounts of noise added to the prior. (**D**) Calibration results on *S.cerevisiae* (GSE144820 [8] only) dataset. Posterior means are cumulatively placed in bins based on their posterior variances. AUPRC for each of these bins is computed against the gold standard (see Methods section for details)

In our comparitive GRN inference analysis, we assess the number of edges predicted in common by each algorithm, on the individual *S. cerevisiae* datasets. We do so by computing the Intersection over Union (IoU) score, filtering each GRN to the top 25% of interactions to remove noisy predictions. Notably, PMF-GRN obtains an IoU score of 15.69%, outperforming alternative algorithms such as SCENIC (3.17%), AMuSR (12.46%), BBSR (14.56%), and StARS (11.78%). The superior performance of PMF-GRN can be attributed to an ability to discern meaningful regulatory interactions, thereby enriching the consensus among predictions. Importantly, our findings underscore a limitation of CellOracle, which achieves an IoU score of 30.28%. This algorithm, while proficient, can only ascertain edges present in the prior-knowledge matrix. Consequently, the two yeast GRNs inferred display high similarity, reflecting an inherent constraint. This characteristic imparts a degree of predictability to CellOracle, limiting its capacity to discover novel interactions beyond the established prior-knowledge. In contrast, PMF-GRNs IoU score is indicative of a more diverse and comprehensive set of common edges. This highlights PMF-GRNs capability to capture nuanced regulatory relationships as a robust and versitle tool for GRN inference.

In a second experiment, we implement a 5-fold cross-validation approach to establish a baseline for each model. Cross-validation is crucial for evaluating the generalization ability of machine learning models like PMF-GRN, particularly in predicting TF-target gene interactions with limited data, a common scenario in experimental settings. To streamline the analysis, we combine the two *S. cerevisiae* single-cell RNA-seq datasets into a single observation matrix. The cross-validation process involves an 80% − 20% split of the gold standard, where a network is inferred using 80% as “prior-known information” and evaluated using the remaining 20%. This process is iterated five times with different random splits to yield meaningful results. We observe that PMF-GRN outperforms SCENIC and CellOracle, while achieving similar performance to BBSR and StARS (Figure 2B). We note that for this experiment, we are unable to implement the AMuSR algorithm as it is a multi-task inference approach that requires more than one task (dataset).

In a third experiment, we evaluate the robustness of each GRN inference method in the presence of noisy prior information. We conduct GRN inference with increasing levels of noise introduced into the prior knowledge. Specifically, the prior information begins with 1% non-zero edges, and we systematically introduce noise to observe the performance of each method. The noise levels are varied from zero noise (original prior, 1% non-zero edges), to 100% noise (resulting in 2% non-zero edges), 250% noise (3.5% non-zero edges), and 500% noise (6% non-zero edges). Our findings, illustrated in Figure 2C, reveal that as the noise in the prior information increases, PMF-GRNs AUPRC experiences a slow decline, mirroring the behavior observed in CellOracle. Notably, PMF-GRN consistently outperforms BBSR, StARS, and SCENIC under these noise conditions, showcasing its robustness in accurately inferring GRNs from noisy priors. These results underscore PMF-GRN as one of the most robust approaches in the face of noisy prior information, thereby emphasizing its utility in practical applications.

To further emphasize PMF-GRN’s robustness in a diverse number of settings, we perform the following two experiments. In the first experiment, we examine the performance of PMF-GRN using different sizes of downsampled yeast expression (Figure 3A). The downsampling procedure involved reducing the expression data to sizes of 80%, 60%, 40%, and 20%, with each size undergoing random sampling five times to generate five distinct datasets per sample size. Remarkably, the AUPRC performance exhibits noteworthy stability across the downsampling variations. Despite the reduction in dataset size, PMF-GRN consistently demonstrates an ability to learn accurate GRNs as evidenced by the sustained AUPRC performance. These findings underscore the robustness of PMF-GRN, suggesting its reliability even under conditions of diminished dataset sizes, a critical consideration for practical applications where data availability may be limited.

**Figure 3:**
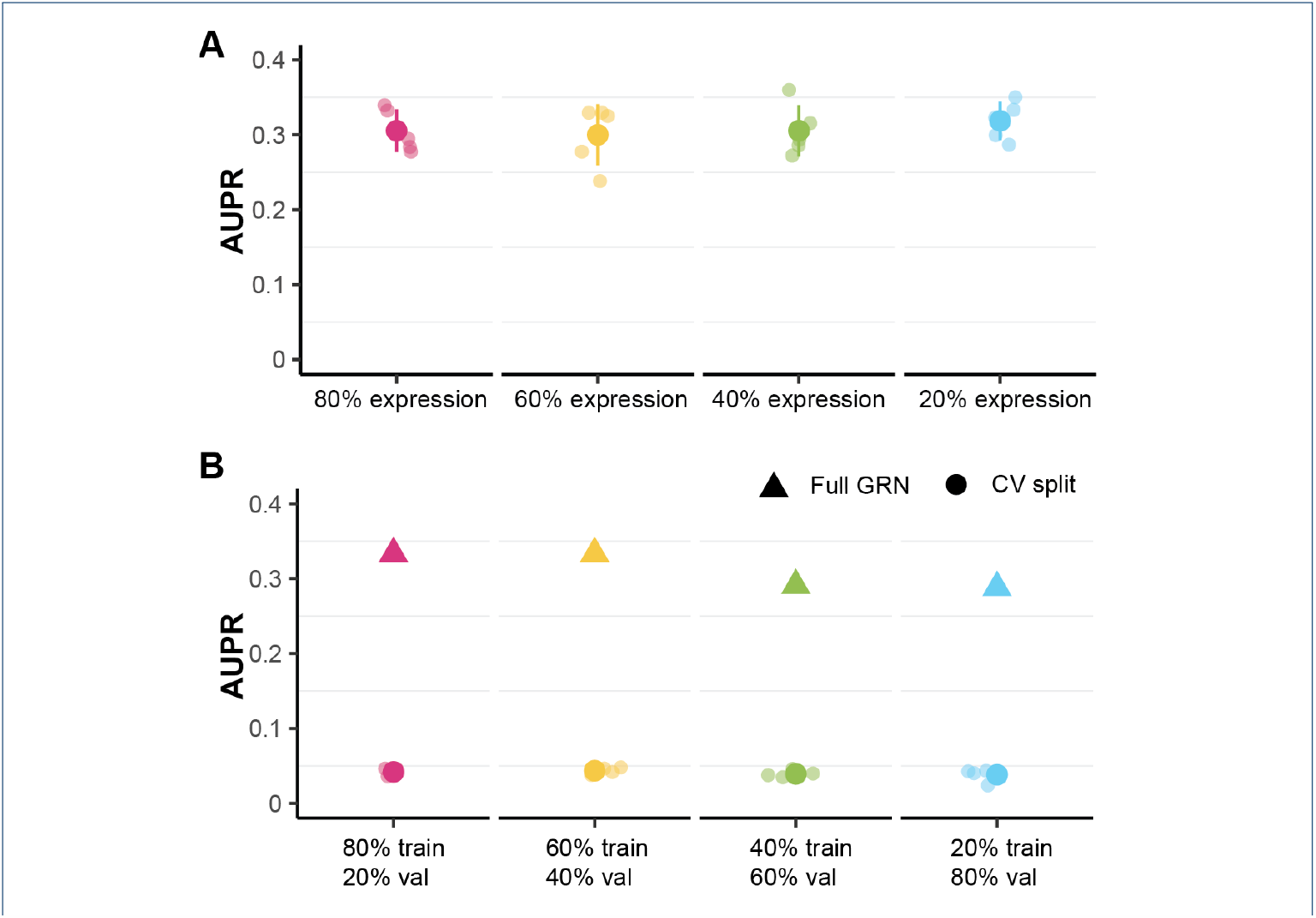
(**A**) GRNs inferred by downsampling *S. cerevisiae* expression data. (**B**) Hyperparameter search performed on 4 different ratios of cross-validation. Dots represent validation AUPRC from hyperparameter search during cross-validation, triangle represents AUPRC from a GRN learned using the most optimal hyperparameters for each ratio.

In a subsequent experiment, we explore the impact of different cross-validation split sizes on hyperparameter tuning for PMF-GRN using the *S. cerevisiae* priorknowledge (Figure 3B). Four distinct cross-validation splits, ranging from 80% training and 20% validation, to 20% training and 80% validation, were employed. For each split, we conducted a hyperparameter search across five samples, selecting the optimal hyperparameters based on the highest validation AUPRC. We then selected the best overall hyperparameters from each split to learn a GRN on the full dataset, in order to demonstrate the downstream effect of cross validation split choice on GRN inference. Surprisingly, our results revealed that the choice of cross-validation split size had a marginal impact on the overall performance of the inferred GRN. Specifically, the AUPRC values for the full GRN remained nearly unchanged regardless of whether an 80% train and 20% validation or 60% train and 40% validation split where employed. Even with more disparate splits, such as 40% train and 60% validation, or 20% train and 80% validation, the decrease in AUPRC was only minor. This implies that PMF-GRN exhibits robustness in hyperparameter selection, with the algorithm consistently converging to optimal settings across varying cross-validation scenarios.

From our experiments on *S. cerevisiae* data, several key observations emerge. First, PMF-GRN consistently outperforms the Inferelator in recovering true GRNs, surpassing two Inferelator algorithms (AMuSR and StARS) and performing similarly to BBSR. Notably, when expression data is not separated into tasks, PMF-GRN outperforms BBSR. In comparison to CellOracle, PMF-GRN demonstrates competitive performance during normal inference and significantly outperforms CellOracle in cross-validation. However, PMF-GRN, in contrast to CellOracle, is not constrained to predicting edges solely within the confines of the prior-knowledge matrix. Furthermore, PMF-GRN consistently outperforms SCENIC across all experiments. A second key observation is that our approach addresses the high variance associated with heuristic model selection among different inference algorithms. When implementing the Inferelator on *S. cerevisiae* datasets under normal conditions, AUPRCs fall within the range of 0.2 to 0.4, showcasing significant variability without a priori information to guide algorithm selection. This diversity among Inferelator algorithms constitutes heuristic model selection, as one cannot predict a priori which algorithm will perform better or discern the reasons behind their divergent performances. In contrast, our method offers reliable results grounded in a principled objective function, delivering competitive performance akin to the best-performing Inferelator algorithm (BBSR) and CellOracle. This underscores the importance of a consistent and robust approach in the face of uncertainty associated with heuristic model selection among disparate algorithms.

To underscore the identifiability issue and affirm the utility of prior-known information, we showcase PMF-GRN’s performance when prior information is unused (e.g., all prior logistic normal means of *A* set to the same low number). This process is replicated for other GRN inference algorithms by providing an empty prior. Additionally, we assess PMF-GRN’s performance when prior-known TF-target gene interaction hyperparameters are randomly shuffled before building the prior distribution for *A*. The results, along with those for the Inferelator and CellOracle, indicate the capability of these approaches to accommodate such prior information effectively.

### PMF-GRN Provides Well-Calibrated Uncertainty Estimates

Through our inference procedure, we obtain a posterior variance for each element of *A*, in addition to the posterior mean. We interpret each variance as a proxy for the uncertainty associated with the corresponding posterior point estimate of the relationship between a TF and a gene. Due to our use of variational inference as the inference procedure, our uncertainty estimates are likely to be underestimates. However, these uncertainty estimates still provide useful information as to the confidence the model places in its point estimate for each interaction. We expect posterior estimates associated with lower variances (uncertainties) to be more reliable than those with higher variances.

In order to determine whether this holds for our posterior estimates, we cumulatively bin the posterior means of *A* according to their variances, from low to high. We then calculate the AUPRC for each bin as shown for the GSE125162 [8] *S*.*cerevisiae* dataset in Figure 2D. We observe that the AUPRC decreases as the posterior variance increases. In other words, inferred interactions associated with lower uncertainty are more likely to be accurate than those associated with higher uncertainty. This is in line with our expectations as the more certain the model is about the degree of existence of a regulatory interaction, the more accurate it is likely to be, indicating that our model is well-calibrated.

### PMF-GRN Integrates Single Cell Multi-Omic Data for GRN and TFA Inference in Human PBMCs

We next evaluate PMF-GRN’s ability to learn informative GRNs in a human cell line by focusing on Peripheral Blood Mononuclear Cells (PBMCs). PBMCs represent an essential component of the human immune system, and consist of Lymphocytes (CD4 and CD8 T cells, B cells, and Natural Killer cells), Monocytes and Dendritic cells. Unraveling the distinct regulatory landscape of PBMCs is an essential task to provide insight into how these immune cells interact, as well as coordinate to maintain homeostasis and respond effectively to infections.

To infer an informative and comprehensive PBMC GRN, we harness information from a large, paired single cell RNA and ATAC-seq multi-omic dataset [42]. We adopt a prior-knowledge matrix of TF-target gene interactions (M genes = 18, 557 by K TFs = 860) as previously constructed by [43] for GRN inference with this multi-omic dataset. In this work, the ATAC-seq data was used as a regulatory mask for ENCODE-derived TF ChIP-seq peaks. Regulatory associations were established through the Inferelator-Prior package based on the proximity of TFs to their potential target genes within 50kb upstream and 2kb downstream of the gene transcription start site. We integrate this prior knowledge with the raw expression profiles of 11, 909 PBMCs from a healthy donor to infer a global PBMC GRN and analyze the TFA profiles of eight annotated cell types and several families of immune TFs within this cell line.

We first investigate whether our predicted TFA clusters into distinct cell-type groups, as annotated by [42]. Using UMAP dimensionality reduction, we are able to determine a near clear distinction between each cell type within PBMCs (Figure 4A). Interestingly, the TFA profiles for each of the T cell sub-types (CD4 T, CD8 T, and other T cells) are closely grouped together, suggesting that these cell types may have a similar lineage or TFA patterns, and may share common transcriptional programs or regulatory networks.

**Figure 4:**
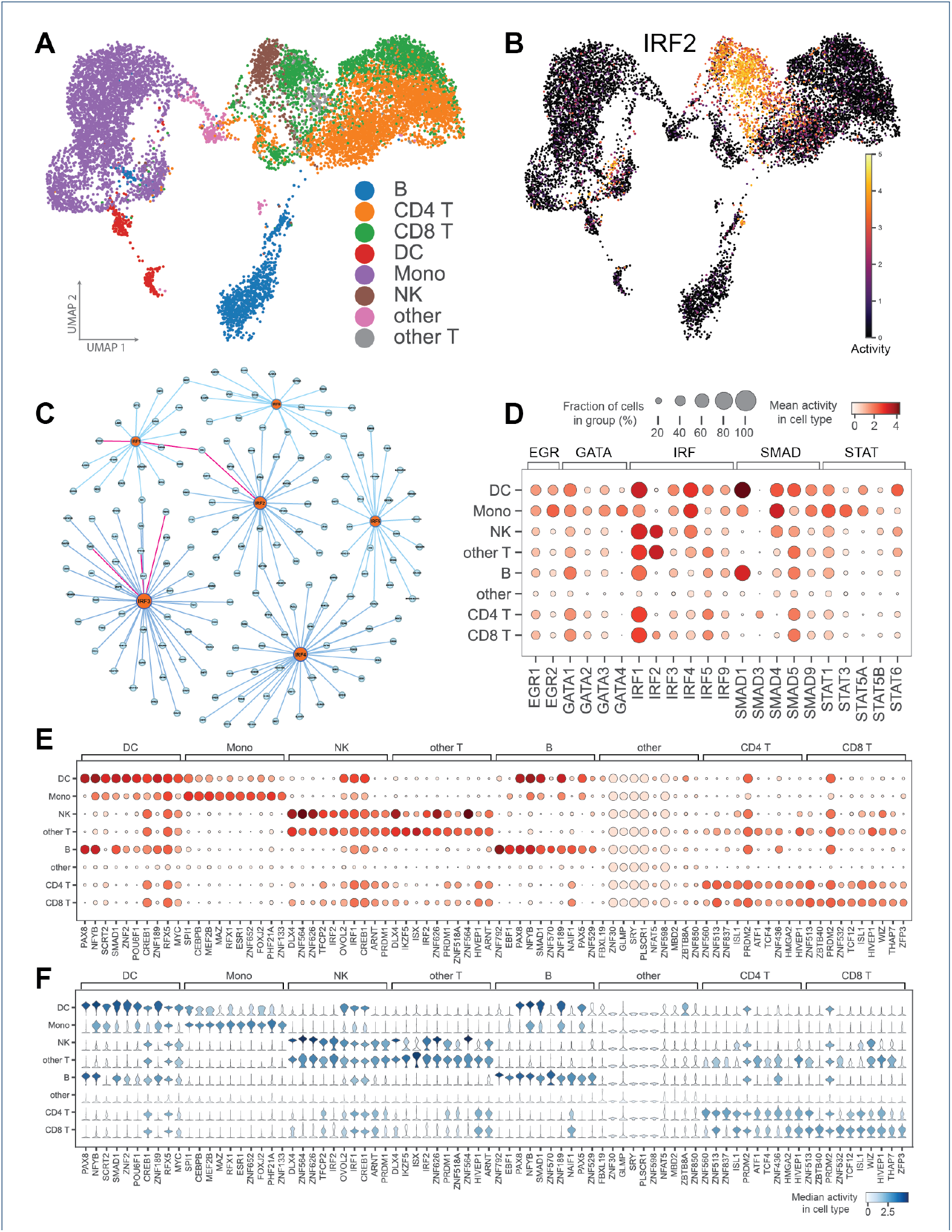
GRN and TFA inference in PBMC. (**A**) UMAP projection of predicted TFA for each annotated PBMC cell type. (**B**) Predicted IRF2 TFA demonstrates high activity in NK and CD8 T cells. (**C**) Heat-map dot-plot depicting TFA of selected immune TFs across annotated PBMC cell types. (**D**) GRN between IRF TFs and their targets. Pink edges indicate literature support for interaction. (**E**) Heat-map dot-plot indicates ten most highly active TFs for each PBMC cell type. (**F**) Violin plot demonstrates corresponding distribution of TFA profiles for ten most highly active TFs.

We next explore the activity profiles of specific immune TF families, starting with the family of TFs belonging to IRF. In PBMCs, IRF contributes to the activation of immune cells that modulate antiviral immunity. Notably, the UMAP projection for IRF2 indicates a high activity pattern within Natural Killer cells and CD8 T cells (Figure 4B). Indeed, IRF2 is essential for the development and maturation of natural killer cells [44], and acts as a CD8 T cell nexus to translate signals from inflammatory tumor microenvironments [45].

In order to support our predicted TFA for the family of IRF TFs, we additionally investigate the regulatory interactions inferred by PMF-GRN (Figure 4C). To do this within a reasonable scale, we first threshold our predicted GRN interactions (described in detail in Methods). Within our thresholded GRN, we predict regulatory edges between IRF1 and the target genes B2M and BTN3A1. IF1 has been documented as a transactivator of B2M [46], while BTN3A1, a defense-related gene, has been found to be upregulated via the IRF1 pathway [47]. Further, we predict that IRF2 also regulates B2M. Supporting evidence demonstrates that IRF2 has been shown to directly bind to genes linked to the interferon response and MHC Class I antigen presentation, including B2M [48]. Finally, we predict regulatory edges between IRF3 and GPR108, RNF5, and TRAF2. GPR108 has been shown to be a regulator of type I interferon responses by targeting IRF3 [49]. Evidence supporting the interaction between IRF3 and RNF5 indicates that RNF5 has an inhibitory effect on the activation of IRF3 [50]. Lastly, TRAF3 has been shown to be a critical component in the activation of IRF3 during the innate immune response to viral infections [51].

In addition to the IRF TFs, several other families of TFs, such as SMAD, STAT, GATA, and EGR, collectively play pivotal roles in PBMCs. These roles contribute to a wide spectrum of functions, including antiviral responses (IRF), fine-tuning immune responses (SMAD), immune cell development (GATA), immediate early responses to signals (EGR), and central regulation of T cells, B cells, and Natural Killer cells (STAT). Their coordinated activities orchestrate the complex interplay of immune cells, enabling PBMCs to effectively respond to diverse stimuli and maintain immune homeostasis.

Similarly to IRF, we also explore edges in our thresholded PBMC GRN for these immune TFs to identify regulatory edges supported by literature. Of the five families of immune TFs that we investigate, we find supporting literature for 60 regulatory edges predicted by PMF-GRN. We provide these literature supported edges, along with their supporting references in Supplemental Table 8. Additionally, we provide a graph representation of each immune TF GRN in Supplemental Figure S2.

We next explore the TFA profiles of each of these immune TFs within the eight PBMC cell types. In Figure 4D, a heat-map dot-plot provides a visual representation of TFA for each immune TF family across the different PBMC cell types. In particular, we observe that within the IRF family, IRF1 is highly active in CD4 T cells. Previous studies have confirmed the pivotal role of IRF1 in CD4+ T cells, where it is essential for promoting the development of TH1 cells through the activation of the Il12rb1 gene [52]. Additionally, SMAD5 is predicted as highly active in B cells. SMAD5 is a key component of the TGF-β signaling pathway, and has been shown to play a crucial role in maintaining immune homeostasis in B cells [53]. We provide a UMAP of the TFA profiles for each of these immune TFs in Supplemental Figure S3.

We further explore our predicted TFA profiles from our global PBMC GRN and calculate the ten most active TFs across the eight distinct cell types. For this experiment, we provide a heat-map dot-plot demonstrating the mean TFA value for each of the top TFs, as well as a corresponding violin plot depicting the distributions of these TFA profiles (Figure 4E and 4F). Visualizing these distinct activity profiles provides a concise and informative snapshot of the predominant TFs contributing significant transcriptional activity within each cell population. For example, within B cells we observe high activity for the TF PAX5. PAX5 is known to play a crucial role in B cell development by guiding the commitment of lymphoid progenitors to the B lymphocyte lineage while simultaneously repressing inappropriate genes and activating B lineage-specific genes [54].

For each annotated cell-type in the PBMC dataset, a set of marker genes were provided. From our ten most active TFs per cell-type analysis combined with their edges to target genes from our thresholded GRN, we find that several of these TFs are predicted to regulate marker genes. For example, within Dendritic cells, the marker gene HLA-DQA1 is predicted to be regulated by the TFs SMAD1 and RFX5; the marker gene HLA-DPA1 is predicted to be regulated by ZNF2 and RFX5; and the marker gene HLA-DRB1 is predicted to be regulated by RFX5. Within CD4 T cells, the marker gene LTB is predicted to be regulated by the TF ZNF436. Within Natural Killer cells, the marker gene PRF1 is predicted to be regulated by ZNF626. Finally, within B cells, the marker gene BANK1 is predicted to be regulated by the TFs ZNF792, EBF1, PAX8, and PAX5; and the marker gene HLA-DQA1 is predicted to be regulated by the TF SMAD1.

From the predicted edges between a snapshot of highly active TFs and annotated marker genes, we find the following supporting evidence. The regulatory relationship between RFX5 and HLA-DQA1 involves the inability of RFX5 to bind to the proximal promoter region of HLA-DQA1, potentially due to DNA methylation, hindering the assembly of active regulatory regions [55]. Additionally, EBF1 orchestrates direct transcriptional regulation of BANK1, leading to the observed downregulation of BANK1 expression [56].

Pairing the intensity (dot-plot) with the distribution (violin plot) of TFA offers a comprehensive view of tje key TFs guiding our regulatory networks. This approach illuminates the variability in their activity levels across diverse immune cell populations, providing a nuanced understanding of the transcriptional dynamics in PBMCs. This information can be used to guide insights into the functional specialization and diversity of immune cells within PBMCs. Further, this comparison provides a sound starting point for exploring the commonalities and differences in the transcriptional regulation of various immune cell populations.

### Evaluating PMF-GRN with BEELINE Synthetic Data

We next evaluated PMF-GRN using synthetic datasets curated from the BEELINE benchmark [37]. This benchmark provides six synthetic networks, linear (LI), linear long (LL), cycle (CY), bifurcating (BF), trifurcating (TF), and bifurcating converging (BFC). In repetitions of ten, expression datasets of increasing cell sizes (e.g., *n* = 100, 200, 500, 2000 and 5000) were generated by sampling. Using these generated expression datasets, as well as the provided reference GRNs, we inferred 300 GRNs using PMF-GRN (Figure 5A). For each of the six synthetic datasets, PMF-GRN outperforms the BEELINE baseline, represented in Figure 5A with a black dashed line.

**Figure 5:**
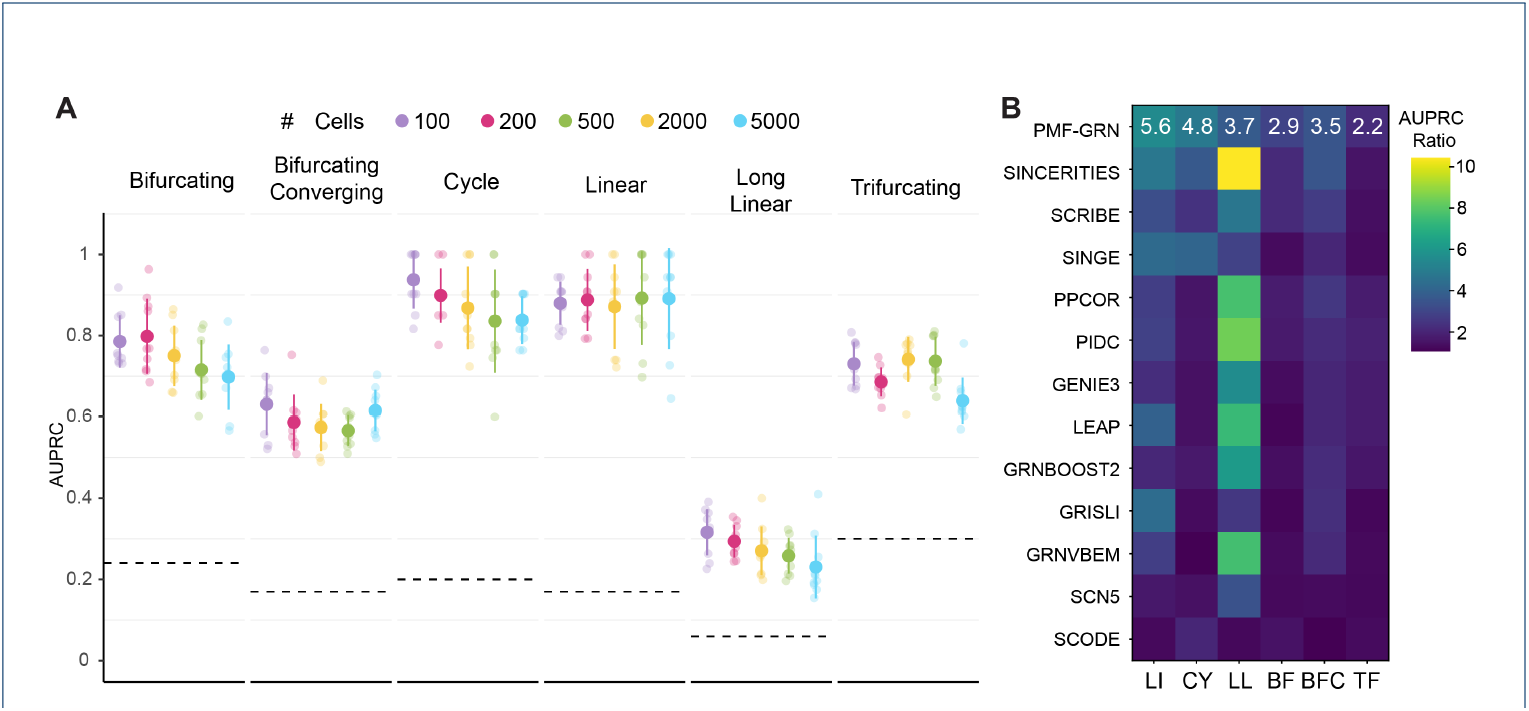
PMF-GRN performance on BEELINE synthetic GRN data (A) PMF-GRN inference performance with half of the ground truth provided as prior network information and the remaining half provided as a gold standard for evaluation. Dashed lines are the expected baseline of a random predictor. (**B**) AUPRC ratio over the baseline random predictor for PMF-GRN in comparison to each of the GRN inference methods used in the original BEELINE benchmark.

To further evaluate PMF-GRN, we calculate the AUPRC ratio of PMF-GRN over the baseline random predictor to compare to the similarly computed ratios in the original BEELINE paper (Figure 5B). We observe that for the linear, cycle, and bifurcating converging, PMF-GRN achieves competitive AUPRC ratios in comparison to the original methods used in the BEELINE benchmark. Interestingly, PMF-GRN does not perform competitively on long linear. This could be due to a number of factors, such as the larger number of intermediate genes introducing additional complexity which PMF-GRN struggles to capture. Alternatively, the extended trajectory introduces a higher-dimensional space, which could present a challenge for our matrix factorization based approach to effectively decompose the data into meaningful latent factors. This presents an interesting avenue of consideration when developing future probabilistic matrix factorization approaches for GRN inference.

## Discussion

In this paper, we introduce a robust framework for probabilistic matrix factorization, optimized through automatic variational inference, to infer GRNs from single-cell gene expression data. A distinctive feature of our approach is the decoupling of the data generation model from the inference procedure, providing unprecedented flexibility. This decoupling allows for modifications to the latent variables and their distributions, without altering the inference process. Such flexibility facilitates the seamless integration of diverse sequencing datasets and modeling assumptions. Unlike previous methods, our framework eliminates the need to define a new inference procedure for each specific dataset or biological context when building new models. PMF-GRN not only offers a flexible and unified approach to GRN inference but also provides a principled methodology for model selection and hyperparameter configuration. The use of a consistent objective function and inference procedure across all generative models streamlines the process of hyperparameter search, reducing ambiguity present in methods like the Inferelator. By conducting hyperparameter search across different generative models, we identify configurations corresponding to optimal values of our objective function, minimizing the reliance on heuristic model selection.

To validate the effectiveness of our approach, we applied PMF-GRN to infer GRNs from single cell *S. cerevisiae* gene expression, comparing results with stateof-the-art single cell GRN inference methods such as the Inferelator, SCENIC and CellOracle. Our method demonstrates competitive, if not superior, performance in terms of AUPRC, in each experiment performed. Here, PMF-GRN provides a stable and reliable inferred GRN without the need for heuristic model selection or data separation into tasks.

Cross-validation experiments further support the robustness of PMF-GRN, BBSR, and StARS, indicating their ability to generalize well to new data without overfitting. In contrast, SCENIC and CellOracle exhibited poor performance during crossvalidation, suggesting potential issues with generalizability. Notably, we assessed the robustness of each algorithm against increasing noise in the prior-knowledge, identifying PMF-GRN and CellOracle as the most resilient to noisy priors. This resilience ensures the reliability of inferred GRNs even in the presence of uncertain prior knowledge.

Our model uniquely provides well-calibrated uncertainty estimates alongside point estimates for each interaction in the final GRN. The evaluation of uncertainty estimates demonstrated that as the posterior variance decreases, the AUPRC increases, indicating that the model is well-calibrated. Biologists can leverage these uncertainty estimates for downstream experimental validation, placing more trust in estimates with lower posterior variance. Finally, the linear scalability of our models computational cost with the number of cells enables its application to single-cell RNA-seq datasets of any size.

Our investigation into PMF-GRN’s application to human PBMCs provides insightful findings into the regulatory landscape of these essential immune cells. Leveraging a comprehensive multi-omic dataset, we demonstrate that our approach integrates single cell RNA and well-curated prior knowledge derived from ATAC-seq data. The resulting global PBMC GRN unveils distinct TFA profiles for eight annotated cell types and various immune TF families. Through UMAP dimensionality reduction, we observe clear clustering of TFA profiles. Focusing on the IRF family, we identify specific TF-target gene interactions supported by literature, shedding light on regulatory relationships critical for immune responses. Extension to other immune TF families reveals their orchestrated activities within PBMCs, contributing to antiviral responses, immune cell development, and the regulation of T cells, B cells, and Natural Killer cells. By exploring predicted edges between active TFs and marker genes, we establish connections between regulatory networks and cellular functions. The combined dot-plot and violin plot visualization strategy provides a nuanced understanding of TF activities, offering a valuable resource for deciphering the intricate transcriptional dynamics in PBMCs. This detailed exploration sets the stage for further investigations into the functional specialization and diversity of immune cells within the PBMC population, with implications for advancing our understanding of immune responses and disease mechanisms.

In the context of synthetic datasets curated from the BEELINE benchmark, PMF-GRN demonstrates robust performance across various network structures. Outperforming the BEELINE baseline across different synthetic networks, PMF-GRN consistently achieves competitive AUPRC ratios compared to the original methods used in the BEELINE benchmark. Notably, PMF-GRN’s competitive performance is observed in linear, cycle, and bifurcating converging structures. However, challenges arise in the long linear structured synthetic data, suggesting potential limitations in capturing the complex dynamics of extended trajectories. Factors such as the increased number of intermediate genes and a higher-dimensional space may contribute to this limitation. This observation opens avenues for future development of probabilistic matrix factorization approaches, encouraging exploration of methods better suited for intricate network structures. The overall success of PMF-GRN in diverse synthetic network scenarios underscores its versatility and effectiveness in inferring GRNs, promising broad applicability in deciphering complex biological systems and regulatory interactions.

## Conclusion

In conclusion, the PMF-GRN framework provides a flexible and principled approach for inferring GRNs from single-cell gene expression data. By decoupling the model and inference procedure, PMF-GRN enables easy integration of new and various sequencing datasets as well as modeling assumptions without the need for defining a new inference procedure. Additionally, PMF-GRN provides a principled approach for model selection through hyperparameter search, reducing the need for heuristic model selection. Overall, PMF-GRN consistently yields high-performing competitive results compared to other state-of-the-art single cell GRN inference methods with a reliable gold standard, and is robust to cross validation, noisy priors and downsampling. Further, PMF-GRN provides well-calibrated uncertainty estimation, enabling a reliable set of results for downstream experimental validation.

We envision many possible directions for future work to design a better algorithm for inferring GRNs under our framework. This framework could be extended to explicitly model multiple expression matrices and their batch effects. We could probabilistically model prior information for *A* obtained from ATAC-seq and TF motif databases, and include this as part of the probabilistic model over which we carry out inference. Evaluating the posterior estimates of the direction of transcriptional regulation, provided by the matrix *B*, could provide a useful benchmark for the computational estimation of TF activation and repression. Research could also be carried out on improved self-supervised objectives for hyperparameter selection.

Future work could also focus on how to use results from our framework to guide experimental wet-lab work. For example, the uncertainty quantification provided by our model could open up new research directions in active learning for GRN inference. Highly ranked, uncertain interactions could be experimentally tested and the results fed back into the prior hyperparameter matrix for *A*. Inference with this updated matrix would ideally yield a better posterior GRN estimate. Posterior estimates of TFA provided by our model could be useful to wet lab scientists, as this quantity provides information about possible post-transcriptional modifications, which are currently challenging to measure experimentally.

Most importantly, the study of GRN inference is far from complete. GRN inference approaches have thus far required new computational models and assumptions in order to keep up with relevant sequencing technologies. It is thus essential to develop a model that can be easily adapted to new biological datasets as they become available, without having to completely re-build each model. We have therefore proposed PMF-GRN as a modular, principled, probabilistic approach that can be easily adapted to both new and different biological data without having to design a new GRN inference method.

## Methods

### Model Details

We index cells, genes and TFs using *n* ∈ {1, · · ·, N }, *m* ∈ {1, · · ·, M } and *k* ∈ {1, · · ·, K}, respectively. We treat each cell’s expression profile *W*_n_ as a random variable, with local latent variables *U*_*n*_ and *d*_*n*_, and global latent variables (that are shared among all cells) σ_*obs*_ and *V* = *A* ⊙ *B*. We use the following likelihood for each of our observations:

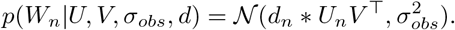

We assume that *U, V*, σ_*obs*_ and d are independent i.e. *p*(*U, V*, σ_*obs*_, *d*) = *p*(*U* )*p*(*V* )*p*(σ_*obs*_)*p*(*d*). In addition to our i.i.d assumption over the rows of *U* and *d*, ee also assume that the entries of *Un* are mutually independent, and that all entries of *A* and *B* are mutually independent. We choose a lognormal distribution for our prior over *U* and a logistic Normal distribution for our prior over *d*:

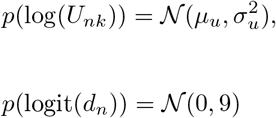

where *μ*_*u*_ ∈ ℝ and *σ*_*u*_ ∈ ℝ ^+^.

We use a logistic Normal distribution for our prior over *A*, a Normal distribution for our prior over *B* and a logistic Normal distribution for our prior over σ_*obs*_:

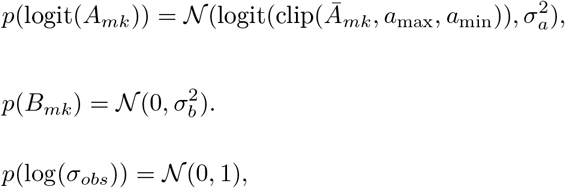

where Ā _*mk*_ ∈ {0, 1}, *a*_max_ ∈ (0, 1), *a*_min_ ∈ (0, 1), *σ*_*a*_ ∈ ℝ _>0_, clip(Ā _*mk*_, *a*_max_, *a*_min_) = max(min(Ā _*mk*_, *a*_max_), *a*_min_) and *σ*_*b*_ ∈ ℝ _>0_. Ā _*mk*_ is given by a pipeline that is used by other methods such as the Inferelator. The pipeline leverages ATAC-seq and TF binding motif data to provide binary initial guesses of gene-TF interactions. *a*_max_ and *a*_min_ are hyperparameters that determine how we clip these binary values before transforming them to the logit space.

For our approximate posterior distribution, we enforce independence as follows:

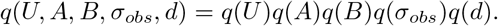

We impose the same independence assumptions on each approximate posterior as we do for its corresponding prior. Specifically, we use the following distributions:

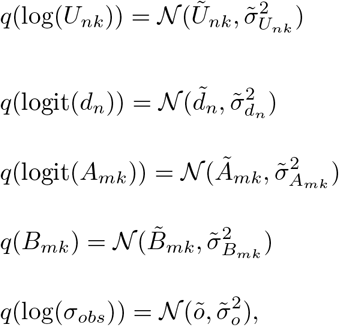

where the parameters on the right hand sides of the equations are called variational parameters; 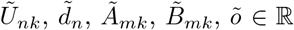and 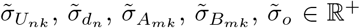 . To avoid numerical issues during optimization, we place constraints on several of these variational parameters.

### Inference

We perform inference on our model by optimizing the variational parameters to maximize the ELBo. In doing so, we minimise the KL-divergence between the true posterior and the variational posterior. In practice, to help with addressing the latent factor identifiability issue, we use a modified version of the ELBo where the prior and posterior terms are weighted by a constant β ≥ 1 [57]:

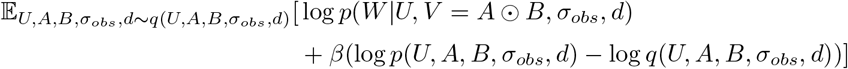

Inference is carried out using the Adam optimizer with learning rate 0.1 and beta values of 0.9 and 0.99. We clip gradient norms at a value of 0.0001. We set *a*_min_ = 0.005, *a*_max_ = 0.995, 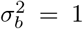and *μ*_*u*_ = 0. We vary *σ*_*a*_ and *σ*_*u*_ as hyperparameters that control the strengths of the priors over *A* and *U*, respectively. We also vary *β* as a hyperparameter.

We choose a hyperparameter configuration using validation AUPRC as the objective function as well as the early stopping metric. We hold out hyperparameters for *p*(A) for a fraction of the genes. We do this by setting Ā _*mk*_ = 0 for m corresponding to these genes for all k. During inference we regularly obtain posterior point estimates for these entries and measure the AUPRC against the original values of these entries as given in the full prior. This quantity is known as the validation AUPRC.

Once we have picked the hyperparameter configuration corresponding to the best validation AUPRC, we perform inference with this model using the full prior without holding out any information. We use an importance weighted estimate of the marginal log likelihood as our early stopping criterion:

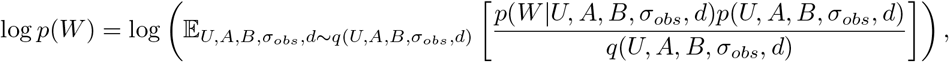

where the expectation is computed using simple Monte Carlo and the logΣ -exp trick is used to avoid numerical issues.

### Computing Summary Statistics for the Posterior

After training the model, we use Ã and 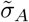, the variational parameters of *q*(*A*), to obtain a mean and a variance for each entry of A. Since *q*(*A*) is logistic normal, it admits no closed form solution for the mean and variance. We therefore use Simple Monte Carlo i.e. we sample each entry of A several times from its posterior distribution and then compute the sample mean and sample variance from these samples. We use each mean as a posterior point estimate of the probability of interaction between a TF and a gene, and its associated variance as a proxy for the uncertainty associated with this estimate.

### Calculating AUPRC

The gold standards for the datasets used in this paper do not necessarily perfectly overlap with the genes and TFs that make up the rows and columns of *A* as defined by the prior hyperparameters i.e. there may be genes and TFs in the gold standard with a recorded interaction or lack of interaction, that do not appear in our model at all because they are not present in the prior. The reverse is also true: the prior may contain genes and TFs that are not in the gold standard. For this reason, we compute the AUPRC using one of two methods: ‘keep all gold standard’ or ‘overlap’, which correspond to evaluating only interactions that are present in the gold standard or only interactions that are present in both the gold standard and the prior/posterior. We present results with ‘keep all gold standard’ AUPRC as the evaluation metric when comparing our model to the Inferelator in Figures 2 and S1. For our evaluation of uncertainty calibration (Figure 2 D), we use the overlap AUPRC so that bins containing a lower number of posterior means do not have artificially deflated AUPRCs (see the Evaluating Calibration of Posterior Uncertainty part of the Methods Section for further information).

### Evaluating Calibration of Posterior Uncertainty

We create 10 bins, corresponding to the lowest 10%, 20%, 30% and so on of posterior variances. We place the posterior point estimates of TF-gene interactions associated with these variances into these bins and then calculate the ‘overlap AUPRC’ for each bin using the corresponding gold standard. The AUPRC for each bin is calculated using those interactions that are in the gold standard and also in the bin. We use such a cumulative binning scheme because using a non-cumulative scheme could result in some bins having very small numbers of posterior interactions that are present in the gold standard, which would lead to noisier estimates of the AUPRC.

### Inference and Evaluation on Multiple Observations of W

The Inferelator method applies two scRNA-seq experiments separately on *S. cerevisiae*, with each resulting in a distinct model. These models are used to infer TF-gene interaction matrices, which are then sparsified. The final matrix is obtained by taking the intersection of the two matrices and retaining only the entries that are non-zero in both matrices. In our approach, we also train a separate model on each expression matrix, and obtain a posterior mean matrix for *A* for each of them. To obtain the final posterior mean matrix for *A*, we average the posterior mean matrices from each model. While this approach works well, future research could focus on explicitly modeling separate expression matrices within the model, as mentioned in the Discussion section.

### Measuring the Impact of Prior Hyperparameters

We evaluate the utility of each of the prior hyperparameter matrices used in our experiments. In Figures 2A and S1A, we present with grey dots the AUPRCs achieved when performing inference using shuffled prior hyperparameters for *A*. This corresponds to randomly assigning to each row (gene) of *A*, the prior hyperparameters that correspond to a different row of A. Shuffling the hyperparameters should lead to worse performance, as the posterior estimates should then also be shuffled, whereas the row/column labels for the posterior will remain unshuffled. For the ‘no prior’setting, shown with black dots in the figures, we set Ā _mk_ = 0 ∀ *m, k*. The difference in AUPRC achieved using the unshuffled vs shuffled or no hyperparameters measures the usefulness of the provided hyperparameters for the inference task on the dataset in question.

### Cross-Validation

For *S. cerevisiae*, we perform a five-fold cross validation experiment (Figure 2B). Cross-validation is performed by partitioning the gold standard into an 80% - 20% split, where 80% of the data represents prior-known information to be used as a prior for *p(A)*, and the remaining 20% is treated as the gold standard for evaluation. This process is repeated five times to generate five random splits of the data in order to robustly evaluate GRN inference. It is important to note that PMF-GRN performs hyperparameter search before inferring a final GRN within each cross-validation split. For each of the five partitioned cross-validation folds the 80%, or prior portion, is further split into 80% train and 20% test for hyperparameter search and evaluation. Once the optimal hyperparameters have been determined, the initial 80% split is treated as the training data, while the remaining 20%, which was not seen during hyperparameter selection, is used for evaluation.

### Intersection over Union

Intersection over Union (IoU) scores were computed using the GRN learned by each algorithm for the two *S. cerevisiae* expression datasets. For each GRN, we calculate and retain the top 25% of predicted edges in order to obtain the best estimates for each algorithm and elimate noisier predictions. For each algorithm, we compute both the intersection and the union of the GRN interactions predicted from the two *S. cerevisiae datasets*. Dividing the Intersection by the Union allows us to obtain a score indicating how similar the two inferred GRNs are for each algorithm.

### Downsampling Expression

For *S. cerevisiae*, in repetitions of five, we randomly sample the *S. cerevisiae* expression matrix on the cell axis to obtain downsampled expression dataset sizes of 80%, 60%, 40%, and 20%. We perform a hyperparameter search, using an 80% training - 20% validation split of the prior-knowledge matrix, on each of these five expression matrices for each sample size. Using these hyperparameters, we infer GRNs for each repetition within each split to obtain our final downsampled GRNs.

### Exploring the Effect of Cross-Validation Ratios on Hyperparamater Selection

To effectively explore the influence of cross-validation split size on obtaining optimal hyperparameters for GRN inference, we methodically separate our S. cerevisiae prior-knowledge into 4 different split sizes. These splits consist of 80% training - 20% validation, 60% training - 40% validation, 40% training - 60% validation, and 20% training - 80% validation. For each split size, we obtain 5 random training and validation splits to ensure robust results. We then perform hyperparameter search across each 5 random splits for each split size. Using the best overall hyperparameters for each split size, we infer a final GRN to demonstrate the impact each particular split had on obtaining the optimal hyperparameters for the final GRN.

### Datasets and Preprocessing

We inferred each GRN using a single-cell RNA-seq expression matrix, a TF-target gene connectivity matrix, and a gold standard for bench-marking purposes. We modeled the single-cell expression matrices based on the raw UMI counts obtained from sequencing for the *S. cerevisiae* and PBMC datasets, which were therefore not normalized for the purpose of this work. For the two *B. subtilis* datasets used in this work, we demonstrate the effect of different normalization and scaling techniques, and convert all data used to integers in order to create a single-cell-like dataset. We further obtained binary TF-gene matrices representing prior-known interactions, which served as prior hyperparameters over A, and were derived from the YEASTRACT and subtiwiki databases, as well as from [43] for PBMC. We acquired a gold standard for *S. cerevisiae* our datasets from independent work which is detailed below.

#### Saccharomyces cerevisiae

We used two raw UMI count expression matrices for the organism *S. cerevisiae* obtained from NCBI GEO (GSE125162 [8] and GSE144820 [40]). For this well studied organism, we employed the YEASTRACT [58, 59] literature derived network of TF-target gene interactions to be used as a prior over A in both *S. cerevisiae* networks. A gold standard for *S. cerevisiae* was additionally obtained from a previously defined network [41] and used for bench-marking our posterior network predictions.We note that the gold standard is roughly a reliable subset of the YEASTRACT prior. Additional interactions in the prior can still be considered to be true but have less supportive evidence than those in the gold standard.

#### Peripheral Blood Mononuclear Cells

We used a paired multi-omic single cell RNA-seq and ATAC-seq dataset for PBMC obtained from [42]. The single-cell expression matrix contained 11,909 cells. The prior-knowledge matrix was constructed using the ATAC-seq data from this multiomic dataset, constructed and described in detail by [43]. The prior-knowledge matrix is 18, 557 genes by 860 TFs, and contains 0.5% non-zero edges.

Due to the complex and dynamic nature of PBMCs, a gold standard is currently unavailable for this cell line. To evaluate our inferred network, we implement a 5-fold cross-validation procedure where our chromatin accessibility-based prior is split into 5 random sets, where 80% is used as prior knowledge and 20% is used as the gold standard for evaluation. We then took the intersection of the regulatory edges inferred across each of the 5 fold cross-validation experiments, and filtered to retain the highest quality edges, obtaining a prediction probability of 90% or higher.

#### BEELINE Synthetic Datasets

We used the BEELINE synthetic expression datasets [37] without modification. Reference GRNs were transformed into cross-tab matrices in order to use this information for prior-knowledge and gold standard evaluation. We used 50% of the reference GRN as the prior and the remaining 50% as the gold standard, as was similarly done in [31].

## Declarations

### Data Availability

The datasets used in this work are publicly available. They are referenced in the Methods section and are available through https://github.com/nyu-dl/pmf-grn.

### Code Availability

Code, inferred GRNs, and inference and evaluation scripts can be found at

https://github.com/nyu-dl/pmf-grn.

## Author Contributions

CSG and KC contributed to Conceptualization of the project. OM and KC designed the probabilistic model. OM implemented PMF-GRN Software, Experiments and Validation. CSG implemented PMF-GRN Experiments, Validation, and Inferelator Software. OM, CSG, and KC contributed to Methodology, Software, Validation, Formal Analysis, Visualization, and Writing Original Draft Preparation. CSG contributed to Data Curation. KC and RB contributed to Supervision, Project Administration and Funding Acquisition.

## Acknowledgements

We thank members of the Bonneau lab for insightful discussions and feedback on this manuscript. We also thank the the staff of the NYU IT High Performance Computing and Flatiron Institute Scientific Computing Core. CSG is grateful to Yanis Bahroun, Daniel Berenberg, and Maggie Beheler-Amass for insightful discussions related to this work. This work was supported by Samsung Advanced Institute of Technology (*under the project Next Generation Deep Learning: From Pattern Recognition to AI* ); NSF Award 1922658 NRT-HDR: FUTURE Foundations, Translation, and Responsibility for Data Science; the National Institutes of Health (RM1HG011014, R01NS116350, R01NS118183, R01AI130945); and the Simons Foundation. This material is based upon work supported by the National Science Foundation Graduate Research Fellowship Program under Grant No. DGE-2234660. Any opinions, findings, and conclusions or recommendations expressed in this material are those of the author(s) and do not necessarily reflect the views of the National Science Foundation.

## Supplementary Information

### Appendix A: Supplementary Experiments

#### PMF-GRN Recovers True Interactions in Prokaryotes as Evaluated by Cross-Validation

To demonstrate GRN inference on a forth additional dataset, we carry out experiments using two microarray datasets for the prokaryote *Bacillus Subtilis* (B1 - GSE27219 [60] and B2 - GSE67023 [61]). Although PMF-GRN is not primarily designed to learn GRNs from microarray data, we show that it is still possible to learn informative GRNs with this data. For our *B. subtilis* experiments, we have access to prior-knowledge derived from the subtiwiki database [62, 63, 64]. Here, we implement a 5 fold cross-validation approach by using five random splits of the subtiwiki database-derived information, where 80% is used as prior knowledge and 20% is used as the gold standard for evaluation.

**Supplemental Figure S1:**
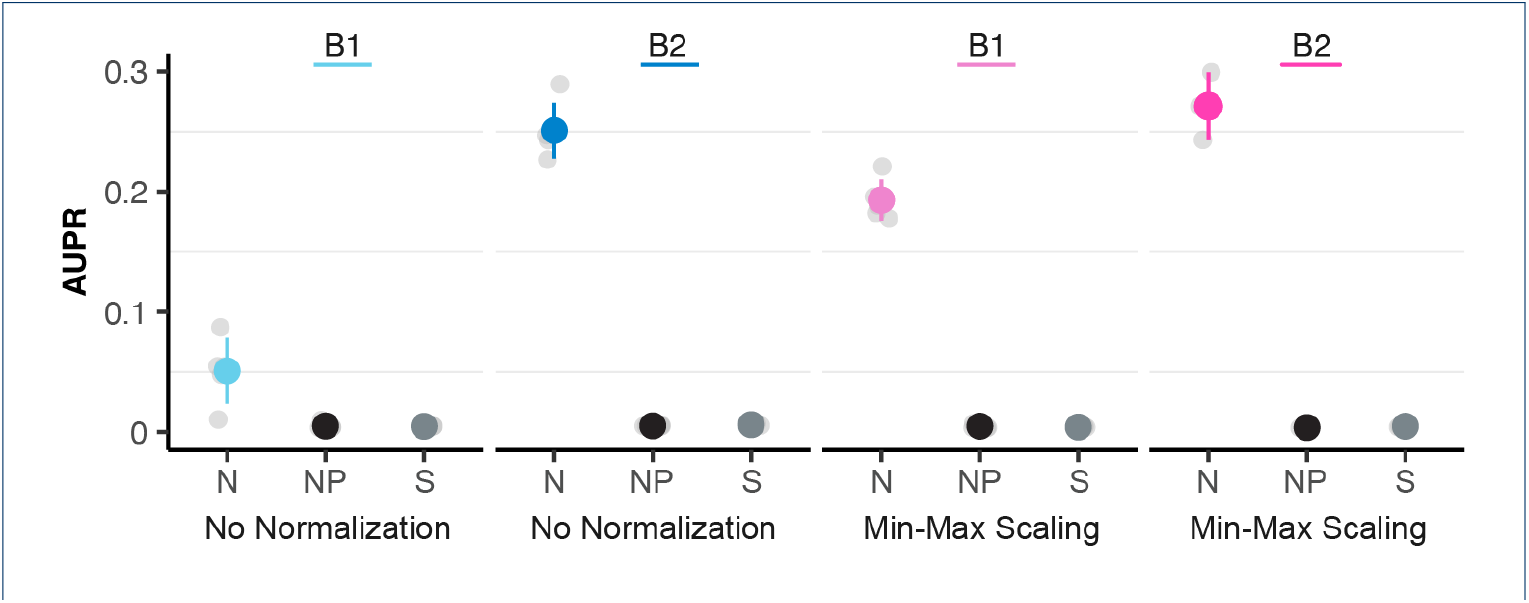
Results for GRNs learned in *B. subtilis* datasets B1 (GSE27219) and B2 (GSE67023) comparing “No Normalization” to “Min-Max Scaling”. Colored dots represent the normal (N) GRN, with the line indicating the mean of the cross-validation experiments ± standard deviation. Negative controls are demonstrated by black dots for GRNs inferred with No Prior (NP) and grey dots for Shuffled Prior (S).

The two *B. subtilis* datasets were previously normalized after data collection as part of standard microarray processing. However, each dataset was normalized using different approaches (described in Methods). For B1, the expression data underwent no further normalization and was simply converted to integers to simulate single-cell-like data. For B2, the expression data was re-scaled and then converted to integers, in order to contain only positive integers resembling single-cell-like data. The results from our experiments are shown in Figure S1, and the numbers used to create this figure are given in Supplementary Table 9 and 10. Using five repeats of cross-validation, we show the performance of GRNs inferred for the two *B. subtilis* datasets (B1 and B2). We remark that the difference in performance between B1 and B2 is likely a result of the chosen microarray processing normalization. To further support this claim, we demonstrate GRN performance after re-scaling the data with min-max scaling (Supplemental Figure S1).

‘No Prior’ and ‘Shuffled’ results are also shown in Figure S1 by black and gray dots respectively. Here, we are able to demonstrate that for B1 and B2, each GRN yields a better performance as compared to negative controls.

## Appendix B: Supplementary Methods

### B.1 TF Target Gene Connectivity Matrix Generation

#### B.1.1 Saccharomyces cerevisiae

Datasets were obtained from [31] without further modification.

### B.1.2 Peripheral Blood Mononuclear Cells

Datasets were obtained from [43] without further modification.

#### B.1.3 BEELINE

Datasets were obtained from [37]. TF-target gene matrices were constructed by creating cross-tab matrices from the RefGRN. 50% of this matrix was used as prior-knowledge, the remaining 50% was used as a gold standard for evaluation.

#### B.1.4 Bacillus subtilis

A prior-known TF-target gene interactions matrix was obtained from the Subtiwiki database [65] from “regulations” (downloaded 07/21/22). Using the columns “regulator locus” and “gene locus” a cross-tab integer matrix was created, where 1 represents the existence of an interaction and 0 represents no interaction. This matrix was randomly split 5 times in 80%-20% proportions along the gene axis to generate independent prior-known information and gold standard matrices.

### B.2 Peripheral Blood Mononuclear Cells

For GRN inference in PBMC, we perform a hyperparameter search using a 5 fold cross-validation split of the prior-knowledge matrix. Here, 80% of the prior-knowledge is used for training, while the remaining 20% is used for validation. For PBMC, we do not have access to a gold standard network. For this reason, to evaluate our inferred GRNs, we select the network for each of the 5 cross-validation splits that achieves the best training hyperparameters. We then take the intersection of each of these 5 optimal networks, and filter the predicted interactions by those obtaining predicted means (probability) of > 0.90 across every split.

For TFA inference, we select the best overall hyperparameters from the 5 fold cross-validation hyperparemeter search to infer a single network. We do this in order to obtain a single TFA matrix where the entries of this matrix are not affected by averaging across multiple datasets. All predicted TFA values are non-zero by nature of matrix factorization. We thus apply an l1 regularization on the matrix with λ = 1 to push low scoring activity values to 0.

UMAP projections were performed on this regularized TFA matrix using the scanpy package [66] with the following parameters: n-neighbors= 10, n-pcs= 40. Additional UMAPs for each of the considered PBMC immune TFs are available in Supplemental Figure S3. To create the GRN diagrams for PBMC, we used the Gephi Open Graph Viz Platform [67]. Additional GRNs for each of the PBMC immune TF families are displayed in Supplemental Figure S2 PBMC heat-map dot-plots and violin-plot were created using the default scanpy parameters for these functions.

**Supplemental Figure S2:**
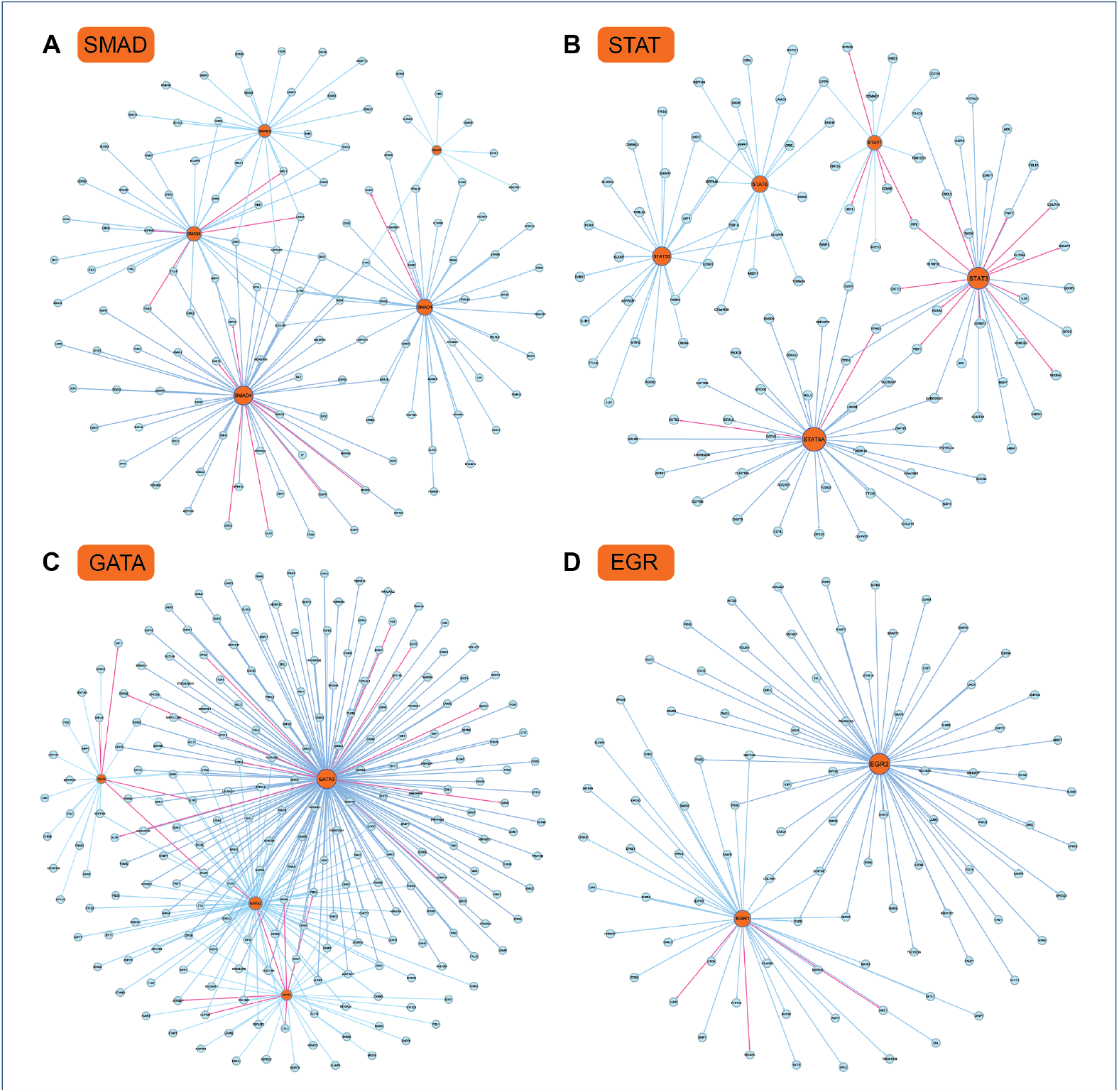
PBMC GRN graphs for the family of TFs belonging to (**A**) SMAD, (**B**) STAT, (**C**) GATA, and (**D**) EGR. TFs for each graph are represented by orange nodes, while target genes are blue nodes. Pink regulatory edges represent interactions with supporting literature. Color scale for blue regulatory edges are scaled from light blue (less out-degree regulation) to darker blue (more out-degree regulation).

**Supplemental Figure S3:**
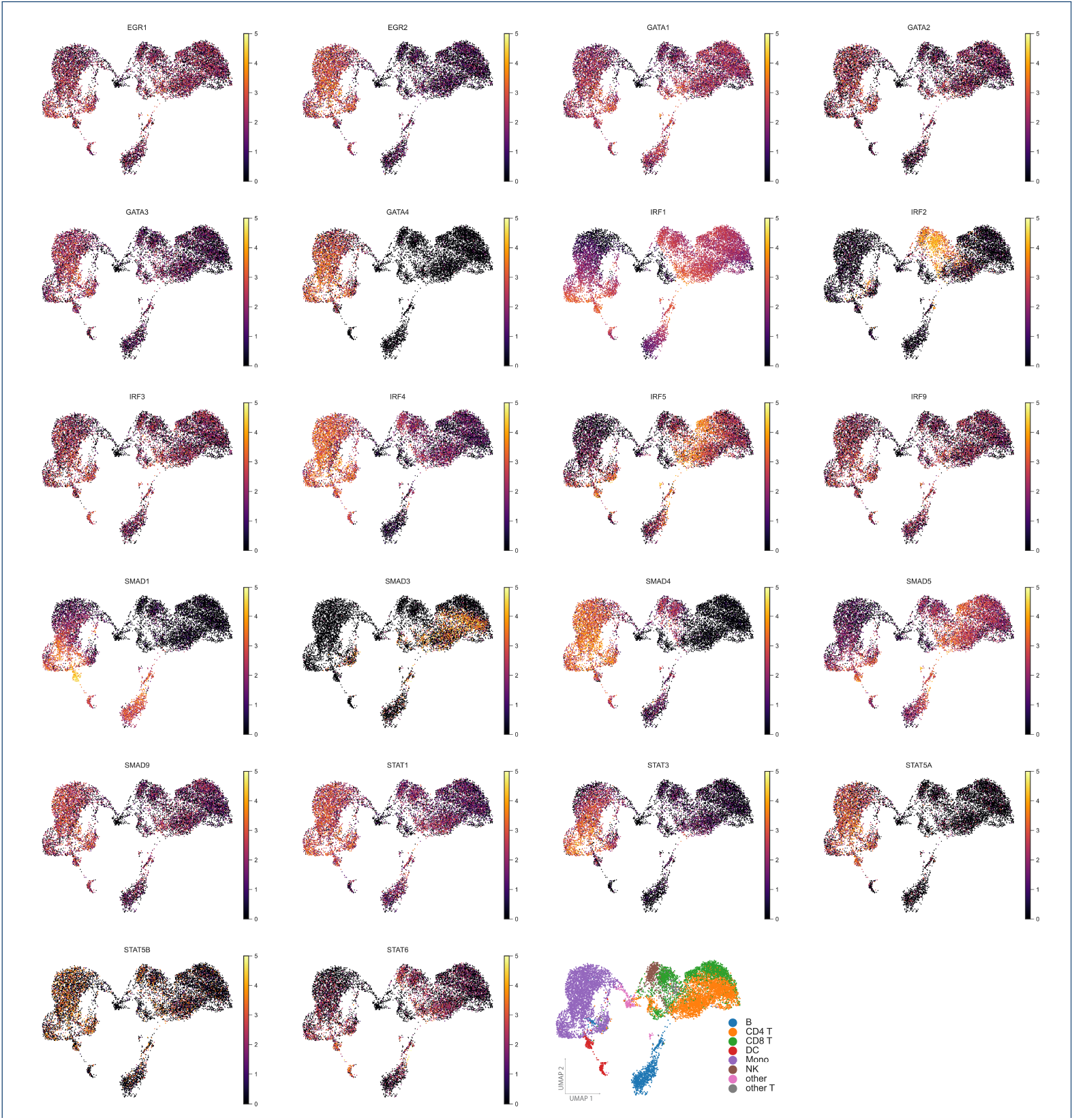
UMAP of predicted PBMC TFA. Each UMAP highlights the specific location of activity for each TF considered from the immune PBMC TFs. The final UMAP serves as a reference to TFA annotated by cell-type.

#### Bacillus subtilis

We used two microarray datasets for *B. subtilis*, which we label as B1 (GSE27219) and B2 (GSE67023). Both B1 and B2 underwent different normalization as part of standard microarray processing, described in detail in [60] and [61]. For the experiment “No Normalization”, B1 was simply converted to integers, while B2 contained negative numbers and had to be scaled and then converted to integers so that the data represented positive integers similar to single-cell data.

To demonstrate the importance of scaling microarray data to place independently collected datasets on the same scale, we demonstrate how Min-Max Scaling improves inference in both *B. subtilis* datasets. For “Min-Max Scaling”, both B1 and the positive scaled B2 dataset were subsequently normalized using the following logic. Using the observation axis, values were linearly transformed so that the minimum value was mapped to 0 and the maximum value was mapped to 1. Each value was then multiplied by 100 and converted to integers to produce the resulting expression matrix of scaled single-cell-like integers.

### B.3 Inferelator, Scenic, and CellOracle Networks

#### B.3.1 Saccharomyces cerevisiae

Networks were inferred using the “multitask” workflow setting of the Inferelator for the same single-cell *S.cerevisiae* datasets described in [31]. For each algorithm, BBSR, StARS, and AMuSR, the following parameters were used: gold_standard_filter_method=“keep_all_gold_standard”, num_bootstraps=5. Aggregated multi-task networks were used for benchmarking, while single-task networks were disregarded for the purpose of this work. To make these networks directly comparable to PMF, we did not make use of normalization, count minimum, or meta-data options available within the Inferelator workflow.

Networks inferred with Scenic and CellOracle used the same input files, with no additional parameters specified.

## Appendix C: Supplementary Tables

**Supplementary Table 1:** GRN inference method comparison table. Table includes input data, output, methodology and pipeline organization for each of the six GRN inference methods discussed in this work.

| Method | Input | Output | Methodology | Pipeline |
| --- | --- | --- | --- | --- |
| PMF-GRN | (1) one or more scRNA-seq datasets<br>(2) priors constructed from genomic data such as ATAC-seq or ChIP-seq and TF motifs or literature database derived priors | (1) individual (and) combined GRN<br>(2) TFA | probabilistic matrix factorization, variational inference | (1) hyperparameter search using 80-20 split of input prior<br>(2) GRN inference with optimal model parameters<br>(3) evaluate GRNs with AUPRC, MCC and F1 scores and uncertainty calibration |
| SCENIC | (1) scRNA-seq<br>(2) a list of TFs ranking databases for motif analysis | (1) GRN: loom file containing regulons (interactions between TFs and their target genes)<br>(2) TFA: biological activity of regulons in a given cell | tree based regression models, gradient boosting machine regression, motif enrichment analysis, and gaussian mixture models for regulon activity binarization | (1-4) preprocessing steps<br>(5) network inference<br>(6) module generation<br>(7) motif enrichment and TF regulon prediction<br>(8) cellular enrichment<br>(9) optional binarization of cellular regulon activity<br>(10) clustering of cells based on regulon activity |
| CellOracle | (1) ATAC-seq + TF motifs<br>(2) scRNA-seq | (1) GRN<br>(2) cell-state transition vectors after gene perturbation | Bayesian Ridge regression | (1) base GRN construction with scATAC-seq or promoter databases<br>(2) scRNA-seq preprocessing<br>(3) context dependent GRN inference<br>(4) network analysis<br>(5) simulation of cell identity after TF perturbation<br>(6) calculation of pseudotime gradient for perturbation scores |
| AMuSR | (1) two or more scRNA-seq datasets<br>(2) one or more priors constructed from genomic data and TF motifs or literature database derived priors | (1) individual and combined GRNs<br>(2) TFA | Regularized linear regression | (1) estimating TFA<br>(2) learning regression parameters<br>(3) model selection with bayesian information criterion<br>(4) evaluate GRNs with AUPRC, MCC, F1 scores |
| BBSR | (1) one or more scRNA-seq datasets<br>(2) one or more priors constructed from genomic data and TF motifs or literature database derived priors | (1) individual (and) combined GRN<br>(2) TFA | Bayesian best subset regression | (1) estimate TFA<br>(2) learn regression parameters<br>(3) model selection<br>(4) evaluate GRNs with AUPRC, MCC, F1 scores |
| StARS | (1) one or more scRNA-seq datasets<br>(2) one or more priors constructed from genomic data and TF motifs or literature database derived priors | (1) individual (and) combined GRN<br>(2) TFA | Least absolute shrinkage and selection operator combined with the Stability Approach to Regularization Selection | (1) estimate TFA<br>(2) learn regression parameters<br>(3) model selection<br>(4) evaluate GRNs with AUPRC, MCC, F1 scores |

**Supplementary Table 2:**
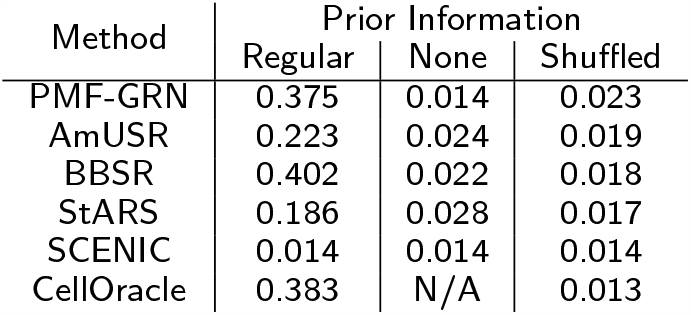
AUPRCs achieved by PMF-GRN, the Inferelator algorithms (AMuSR, BBSR, and StARS), Scenic and CellOracle on S. cerevisiae datasets.

**Supplementary Table 3:** AUPRCs achieved by PMF-GRN, the Inferelator algorithms (BBSR, and StARS), Scenic and CellOracle on S. cerevisiae datasets using the gold standard for 5-fold cross validation.

| Method | Cross Validation Split |  |  |  |  |
| --- | --- | --- | --- | --- | --- |
|  | Split 1 | Split 2 | Split 3 | Split 4 | Split 5 |
| PMF-GRN | 0.114 | 0.096 | 0.086 | 0.1342 | 0.118 |
| BBSR | 0.112 | 0.128 | 0.161 | 0.171 | 0.139 |
| StARS | 0.109 | 0.137 | 0.154 | 0.195 | 0.151 |
| SCENIC | 0.020 | 0.021 | 0.018 | 0.025 | 0.021 |
| CellOracle | 0.034 | 0.042 | 0.034 | 0.043 | 0.034 |

**Supplementary Table 4:** AUPRCs achieved by PMF-GRN, the Inferelator algorithms (BBSR, and StARS), Scenic and CellOracle on S. cerevisiae datasets using increasing amounts of noise added to the prior-knowledge data.

| Method | Noise Added |  |  |  |
| --- | --- | --- | --- | --- |
|  | No Noise | 100% Noise | 250% Noise | 500% Noise |
| PMF-GRN | 0.343 | 0.280 | 0.198 | 0.149 |
| BBSR | 0.264 | 0.208 | 0.186 | 0.174 |
| StARS | 0.136 | 0.125 | 0.114 | 0.118 |
| SCENIC | 0.075 | 0.068 | 0.059 | 0.055 |
| CellOracle | 0.417 | 0.306 | 0.226 | 0.175 |

**Supplementary Table 5:** Intersection over Union (IoU) scores achieved by PMF-GRN, the Inferelator algorithms (AMuSR, BBSR, and StARS), Scenic and CellOracle for GRNs learned on individual S. cerevisiae datasets.

| Method | Intersection over Union (IoU) |
| --- | --- |
| PMF-GRN | 15.69% |
| BBSR | 14.56% |
| AMuSR | 12.46% |
| StARS | 11.78% |
| SCENIC | 3.17% |
| CellOracle | 30.28% |

**Supplementary Table 6:** AUPRCs achieved by PMF-GRN across 4 different downsample sizes (80%, 60%, 40%, and 20%), across 5 samples for each downsample size.

| Expression size (%) | Sample Number |  |  |  |  |
| --- | --- | --- | --- | --- | --- |
|  | 1 | 2 | 3 | 4 | 5 |
| 80% | 0.2838 | 0.2947 | 0.3395 | 0.3325 | 0.2774 |
| 60% | 0.2382 | 0.3295 | 0.3252 | 0.2775 | 0.3296 |
| 40% | 0.3157 | 0.2723 | 0.3599 | 0.2858 | 0.2939 |
| 20% | 0.2995 | 0.3503 | 0.3229 | 0.2868 | 0.3335 |

**Supplementary Table 7:**
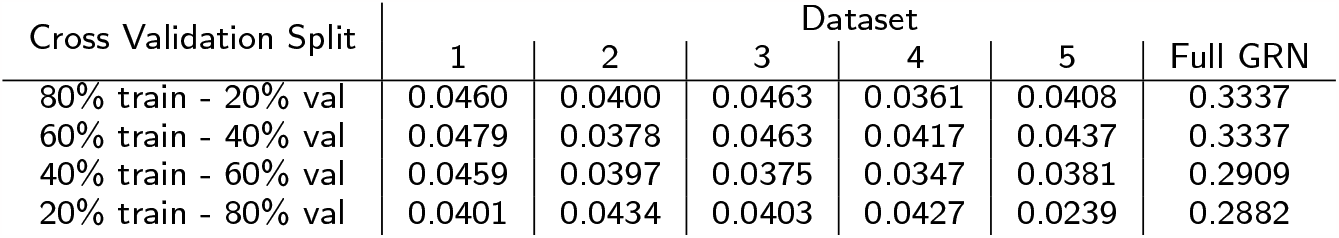
AUPRCs achieved by PMF-GRN across 4 different crossvalidation splits. 5 hyperparameters searches were performed for each cross-validation split. Full GRN was inferred using the hyperparameters for the best overall AUPRC per cross-validation split.

**Supplementary Table 8:**
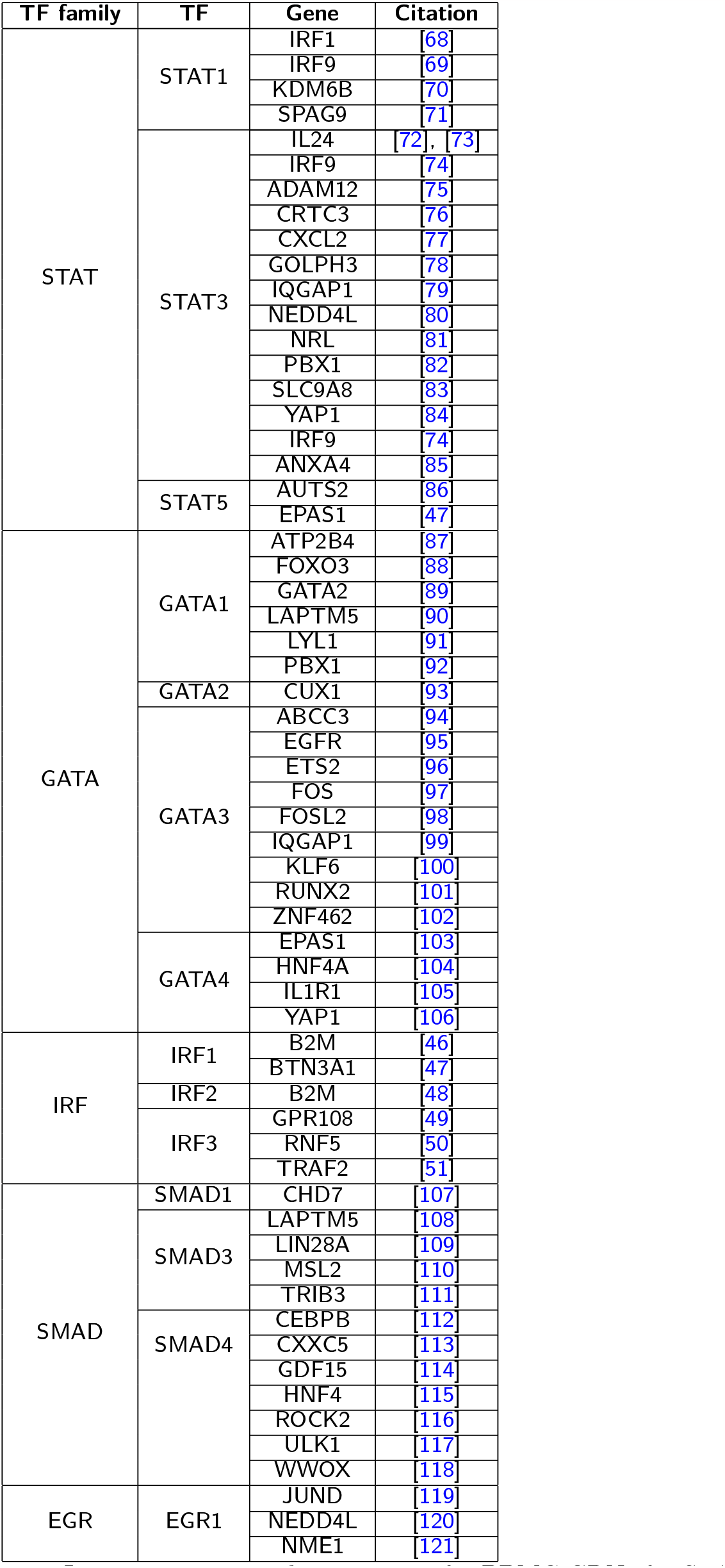
Literature supported interactions for PBMC GRNs for STAT, GATA, IRF, SMAD and EGR immune TF families

**Supplementary Table 9:** AUPRCs achieved by PMF-GRN on *B. subtilis* B1 dataset. Results are reported as the mean AUPRC across five ‘cross-validation’ splits ± standard deviation

| Method | *B. subtilis* Cross-Validation Dataset B1 |  |  |
| --- | --- | --- | --- |
|  | Regular | No Prior | Shuffled |
| No Normalization | $0.0509 \pm 0.0273$ | $0.0048 \pm 0.0003$ | $0.0050 \pm 0.0028$ |
| Min-Max Scaling | $0.1931 \pm 0.0171$ | $0.0042 \pm 0.0003$ | $0.0042 \pm 0.0018$ |

**Supplementary Table 10:** AUPRCs achieved by PMF-GRN on *B. subtilis* B2 dataset. Results are reported as the mean AUPRC across five ‘cross-validation’ splits ± standard deviation

| Method | *B. subtilis* Cross-Validation Dataset B2 |  |  |
| --- | --- | --- | --- |
|  | Regular | No Prior | Shuffled |
| No Normalization | $0.2508 \pm 0.0232$ | $0.0062 \pm 0.0006$ | $0.0052 \pm 0.0004$ |
| Min-Max Scaling | $0.2886 \pm 0.0312$ | $0.0048 \pm 0.0008$ | $0.0038 \pm 0.0005$ |

